# Identification of Multi-landscape and Cell Interactions in the Tumor Microenvironment through High-Coverage Single-Cell Sequencing

**DOI:** 10.1101/2024.06.18.599463

**Authors:** Wenlong Zhong, Ligang Wang, Chengjunyu Zhang, Tonglei Guo, Lihua Zhao, Daqin Wu, Fei Xie, Xiao Wang, Xiuxin Li, Fangxiao Wang, Minghui Li, Weiyue Gu, Tianxin Lin, Xu Chen

## Abstract

Single-cell RNA sequencing (scRNA-seq) is a widely used method for classifying cell types and states and revealing disease mechanisms. However, most contemporary scRNA-seq platforms fail to explore the multilandscape of RNA. Here, we designed a microfluidic chip that combines oligo-dT primers and Random Bridging Co-labeling (RBCL) RNA sequencing to develop an innovative Chigene scRNA-seq technology that can identify gene expression, mutations, and RNA splicing landscapes at the single-cell level. The Chigene scRNA-seq platform demonstrated exceptional performance, with minimal doublet rates of 0.94% (Chigene V1) and 1.93% (Chigene V2). Both versions exhibit high sensitivity, with Chigene V2 achieving nearly 100% RNA coverage and detecting over 1800 genes per cell on average. Targeted capture of single-cell gene mutations enhances mutation detection sensitivity. Moreover, this Chigene V2 platform has been validated in clinical samples for its ability to detect mutations, gene fusions and alternative splicing. The reliability of the platform was further corroborated via known functional gene mutation (CDKN1A) and fusion (FGFR3-TACC). To validate this method’s potential for discovering novel gene mutations in clinical samples, our investigation revealed an intriguing cell subpopulation carrying an ARHGAP5 mutation in urothelial carcinoma. These cells exhibited high-frequency mRNA splicing and exhibited specific crosstalk with T cells, distinguishing them from the subpopulation with the ARHGAP5 wild-type phenotype. Overall, this method provides a robust scRNA-seq platform suitable for comprehensive analyses of clinical specimens at different genetic information levels, thereby offering significant potential in the discovery of novel genes and interactions at the single-cell level.

## Introduction

Single-cell RNA sequencing (scRNA-seq) has revolutionized the field of tumor research by enabling the identification and characterization of cell types, states, lineages, and circuitry at the single-cell level^[1–3]^. These findings have greatly enhanced our understanding of tumor heterogeneity and the immune microenvironment. Genetic mutations^[4]^, gene fusions, and alternative RNA splicing^[5]^ are critical factors that contribute to tumorigenesis and tumor progression. These molecular alterations can significantly impact cellular processes, including cell proliferation, differentiation, metastasis and treatment response^[6–9]^. Integrating transcriptomic analysis with gene mutation data from whole exome sequencing (WES) and RNA alternative splicing data at the single-cell level holds immense potential for advancing the understanding of tumors. Although state-of-the-art scRNA-seq methods are highly sensitive and accurate for quantifying and characterizing cell states^[10–13]^, most of these methods rely on barcoded oligo-dT primers that hybridize to polyadenylated transcripts for RNA capture and cDNA synthesis. This approach primarily detects short fragments located near the poly(A) tail or the 5’ end of the transcript, leaving the remaining sequences of polyadenylated RNA molecules undetected. As a result, the accurate detection of single nucleotide variants (SNVs), gene fusions and alternative splicing events is limited^[14–17]^. Recently, several high-throughput scRNA-seq platforms for total-transcriptome full-length sequencing have emerged ^[18–20]^. However, these methods pose limitations in terms of sequencing depth and mRNA detection sensitivity when total RNA is targeted within a fixed amount of data.

To address the challenges and fulfil the needs of scientific research and clinical practice, we developed a microfluidics-based high-sensitivity and full-length mRNA sequencing method termed Chigene-seq. For Chigene-seq, we employed oligo-dT primers for specific capture of mRNA during reverse transcription, followed by synthesis and specific labelling of the second strand via bridge oligo sequences on the first strand. Subsequently, amplification and sequencing were performed. Additionally, we designed targeted capture steps for specific mutations and gene fusions to further improve the detection efficiency in single cells. Furthermore, we developed a simplified and cost-effective 3’ sequencing approach by using oligo-dT primers only. To validate the performance and clinical applicability of our single-cell sequencing platform, multiple cell lines and human urological cancer samples were utilized. The results indicated the robustness of our platform and allowed for the exploration of gene mutations, gene fusions, and alternative splicing at the single-cell level. Moreover, the application value of this approach was demonstrated by the discovery of novel functional gene mutations in clinical samples. In summary, our novel Chigene scRNA-seq platform offers potential advantages for specific applications, such as mutation detection and alternative splicing analysis, providing expanded methods for discovering new functional genes in clinical samples.

## Results

### Overview of the Microfluidic Chip-based Random Bridging Co-labelling RNA Sequencing Method for Full-length Coverage

We developed a versatile workflow that enables the amplification and fragmentation of mRNA via microbeads carrying oligo-dT primers and random bridging primers on a microfluidic platform (Figure S1). This approach allows the detection of various natural barcodes, including somatic mutations, fusion genes, and RNA isoforms. Based on whether random bridging primers were utilized, the workflow was further divided into two versions: Chigene V1 (conventional 3’ end sequencing) (Figure 1A-B) and V2 (mRNA full-length transcriptome sequencing) (Figure 1C). For Chigene V2, to further increase the detection sensitivity of gene mutation and fusion in single cells, we employed a targeted mutation library enrichment strategy with specific primers. In brief, the core process of this technology is illustrated in Figure 1D. First, the mRNA from single cells is captured on oligo microbeads that contain a ’cellular barcode sequence + polyT’ structure, utilizing the polyA tail of the mRNA. Then the full-length first-strand cDNA was synthesized by reverse transcription. Subsequently, we added a fixed sequence to the end of cellular barcode oligos on the microbeads (as indicated by the three green stars in the legend of Figure 1D) which was unbound mRNA, allowing them to anneal with the bridge oligos that contained random sequences. These bridge oligos serve dual functions: the 5’ end binds and ligates to the modified cellular barcode oligo through complementary pairing, while the 3’ end anneals to the first-strand cDNA at multiple random regions via the random N sequence. Under the action of DNA polymerase, the synthesis of the second-strand DNA is anchored by the bridge oligos, ultimately leading to the systematic incorporation of identical cellular barcode sequences at multiple positions within the cDNA molecules, thereby constructing a dual-strand cDNA library that features systematic single-cell labeling characteristics. This random multipoint bridging strategy locks in cellular origin information at the single-molecule level through the formation of covalent bonds. The bridged and extended products were then amplified to generate sufficient cDNA for library construction and subsequent next-generation sequencing (Figure 1C). To integrate the single-cell mRNA full-length gene expression profiles with the genomic features of interest, we called the molecular features of interest on the basis of genomic features detected by whole exome sequencing (WES) from the same sample. Specifically, we enriched the library for the mutant and fusion genes of interest from the transcriptome library via PCR with specific primers (Figure 1C). By extracting cell barcode and transcript information from the sequencing data, we improved the detection rate of lowly expressed gene mutations. This allowed us to explore the biological functions of cell subpopulations with specific genomic features and their interactions with surrounding cells.

**Figure 1.**
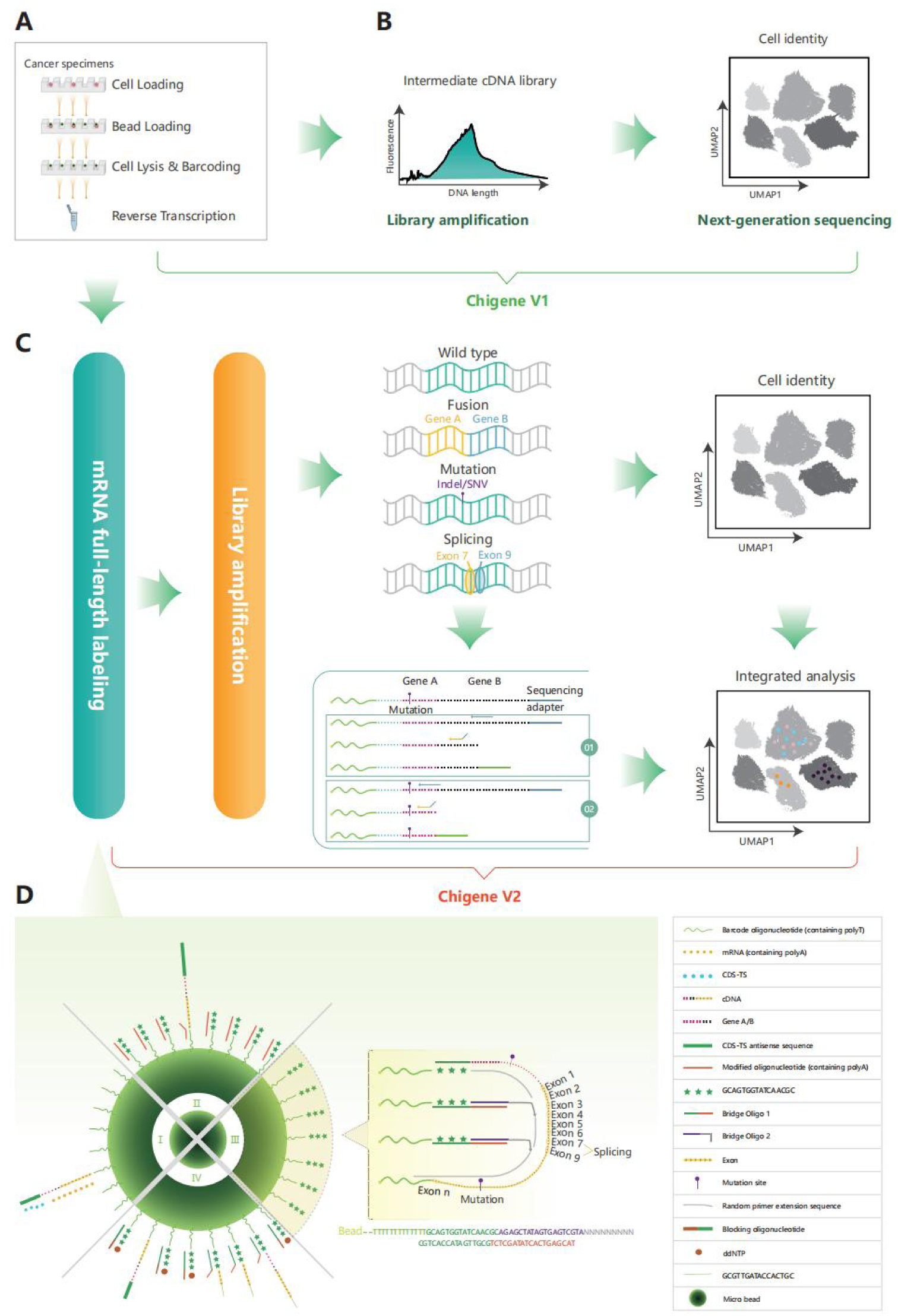
Schematic workflow of Chigene-seq based on the microfluidic chip platform. **(A)** Unique mRNA capture was performed via oligo-dT on a microfluidic chip, and the recovered microbeads were subjected to reverse transcription. The term “cancer specimens” refers to tumor tissue samples obtained from patients diagnosed with urological cancer and collected through biopsy procedures. **(B)** An intermediate cDNA library was amplified and subjected to next-generation sequencing for conventional 3’ scRNA-seq (Chigene V1). **(C)** Overview of the Chigene V2 Library construction and analysis. The process begins with full-length mRNA labelling, followed by library amplification. Different forms of genetic variation, including wild type, fusion (gene A and gene B), mutation (indels/SNVs), and splicing (exon 7 and exon 9), are depicted to highlight the library’s ability to detect these events. The output is used for cell identity determination and integrated analysis. The panel on the middle of Figure 1C provides a detailed view of the sequencing output, indicating the relationships between genes, mutations, and the corresponding sequencing adapters. Panel 01 captures fusion events, whereas Panel 02 captures mutations. **(D)** Detailed schematic illustrating the rationale behind Chigene V2. The circular layout presents the full-length mRNA labeling framework. Notably, the three green stars represent the fixed sequence at the end of the cellular barcode oligos on the microbeads. The legend on the right defines the various components of the schematic.

### Development and validation of the Chigene V1 platform on a microfluidic chip

To evaluate the performance of the microfluidic chip for single-cell sequencing, the Chigene V1 platform, which relies solely on chips and poly T primers, was first developed. To assess the cell capture efficiency and doublet rate of the microfluidic chip, different quantities of HeLa cells ranging from 5000 to 50000 cells were loaded. Microscopic counting revealed that the cell capture efficiency was generally greater than 50%, and the doublet rate was mostly less than 5% (Figure 2A-B). Observations of the microbeads after loading revealed micropores with both single cells and beads (Figure 2C). The time required for cross-contamination between micropores is approximately 5 minutes, whereas cell lysis within the chip is achieved in approximately 12 seconds^[21]^. After the microbeads were loaded, cell lysis, RNA capture, and two quick washes were performed within two minutes to minimize cross-contamination. The recovery of a large portion of the microbeads (∼70%) was achieved (Figure 2D). To evaluate the fidelity of the Chigene V1 platform, a standard mixed-species experiment was conducted using cultured human (A549) and mouse (MC38) cell lines. A mixture of 2500 A549 cells and 2500 MC38 cells was profiled via Chigene V1-seq. The analysis identified 2239 high-quality unique cell barcodes, indicating clear separation of true cells from background noise (Figure 2E). The doublet rate was low at 0.94% (Figure 2F), which is lower than that reported for other scRNA-seq methods (1.6--3.1%), such as Seq-Well^[22]^ and VASAdrop^[19]^. A more detailed comparison of doublet rates with other single-cell platforms can be found in Supplementary Table 2. Additionally, high species specificity of UMI (∼97%) was observed, indicating that high-fidelity single-cell libraries were produced via the Chigene V1 platform (Figure 2G). UMI and gene count distributions revealed that the Chigene V1 platform captured a median of 20,339 UMIs and 4672 genes in single A549 cells, with an average of ∼38k reads, and 17256 UMIs and 4805 genes in single MC38 cells, with an average of ∼30k reads (Figure 2H-I). These results demonstrated the high sensitivity of the Chigene V1 platform.

**Figure 2.**
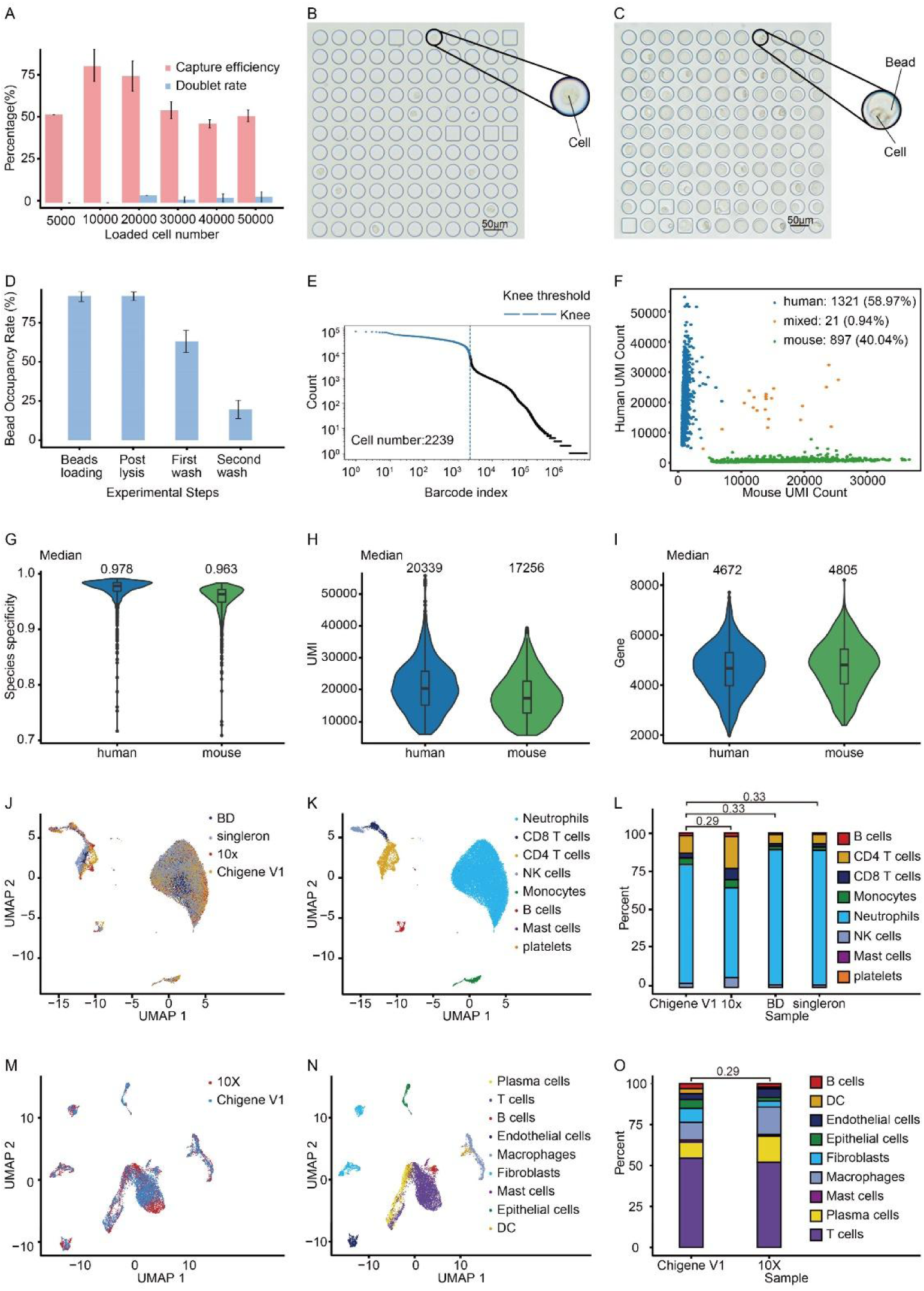
Development and validation of the Chigene V1 platform based on the microfluidic chip. **(A)** Capture rates and doublet rates for various numbers of loaded cells. **(B)** Bright-field images depicting a region of the device after cell loading. **(C)** Bright-field images showing a cell/bead-loaded device. **(D)** Bead occupancy rate following bead loading, cell lysis, and two rounds of washing. **(E)** Barcode plot used for identifying barcodes representing true cells (blue line). The barcodes of the A549-MC38 mixed cells were arranged in descending order on the basis of gene count. **(F)** Species-mixing scatter plot showing the single-cell capture efficiency and doublet rate of Chigene V1-seq. **(G)** Species specificity of UMIs in a human‒mouse cell mixture. The median species specificity of UMIs in human cells (A549) was 0.978. The median species specificity of UMIs in mouse cells (MC38) was 0.963. **(H-I)** Violin plots and box plots showing the number of UMIs and genes detected in individual A549 and MC38 cells. A549 cells: n = 1321; MC38 cells: n = 897. The data depicted in the box plots correspond to the first and third quartiles (lower and upper hinges) and the median (centre). **(J and K)** UMAPs displaying single-cell profiles (dots) from PBMCs, with colors indicating the different platforms used (J) and each cell type (K). **(L)** Proportion of cells of each cell type (y-axis) detected via different platforms (x-axis) in the PBMC libraries. **(M and N)** UMAPs displaying single-cell profiles (dots) of human urachal carcinoma cells, with colors indicating the different platforms used (M) and each cell type (N). **(O)** Proportion of cells of each cell type (y-axis) detected via Chigene V1 and 10x Genomics (x-axis) in human urachal carcinoma libraries.

Next, we evaluated the ability of the Chigene V1 platform to distinguish cell populations in complex primary samples. This was achieved by comparing its performance with that of 10x Genomics, BD Rhapsody, and Singleron using human peripheral blood cells following erythrocyte lysis. Compared with these mainstream scRNA platforms, Chigene V1-seq exhibited very high cluster consistency (Figure 2J and Figure S2A-D) and similar cell proportions (Figure 2K-L and Figure S2E). However, the performance of the Chigene V1 platform was more similar to that of the BD Rhapsody and Singleron platforms because the same cell capture strategy (microfluidic chip) was used. Furthermore, when a more complex urachal carcinoma sample was used for performance comparison between Chigene V1 and 10x Genomics, similar results were observed (Figure 2M-O, Figure S2F-H).

### Development of the Chigene V2 platform based on the Chigene V1 platform and Random Bridging Co-labeling technique

To enhance the capture of full-length mRNAs, we integrated the Random Bridging Co- labelling (RBCL) technique into the Chigene V1 platform and developed the Chigene V2 platform. The gene body coverage of the Chigene V2 platform was substantially greater than that of the Chigene V1 and 10x platforms (Figure 3A). Moreover, species-mixing experiments were conducted to analyse specificity on the Chigene V2 platform. The results revealed acceptable doublet rates (1.93%) and specificity (∼95%) in comparison to those of the Chigene V1 platform (Figure 3B-C). Owing to its ability to capture full-length genes, compared with the Chigene V1 platform, the Chigene V2 platform resulted in a decrease in average UMIs and gene counts per cell at comparable sequencing depths (∼36k reads per human cell and ∼26k reads per mouse cell) (Figure 3D-E). Additionally, the Chigene V2 platform demonstrated more homogeneous coverage of gene body regions than did other high-throughput full-length single-cell RNA sequencing platforms, such as VASA-seq, snRandom, and bulk RNA-seq (Figure 3F). Compared with the canonical full-length scRNA-seq assays (Smart-seq2), Chigene V2 exhibited similar gene body coverage uniformity (Figure S3A). Gene body coverage analysis of the GAPDH and ACTB genes revealed that unlike poly(A)-based 10x Genomics, which exhibited pronounced bias toward the 3’- end, Chigene V2, VASA-seq, and snRandom did not display significant 3’- or 5’-end bias across the gene body. As expected, Chigene V2 displayed greater coverage depth, which was attributable to its emphasis on capturing mRNA within a comparable sequencing data volume (Figure 3G-H).

**Figure 3.**
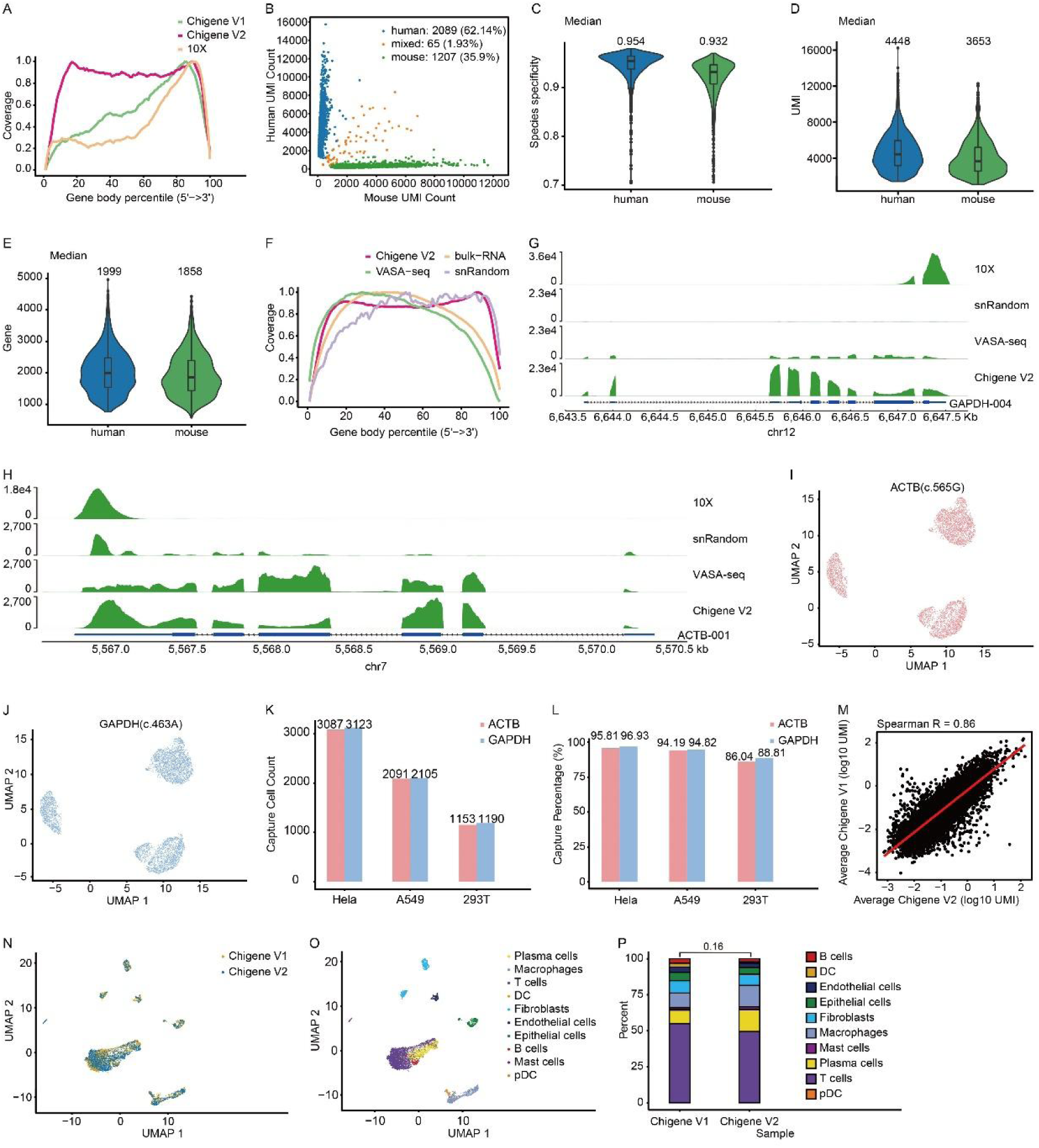
Development of the Chigene V2 platform based on the Chigene V1 platform and Random Bridging Co-labeling technique. **(A)** Gene body coverage shown by read distribution in three platforms. **(B)** Scatter plot illustrating the single-cell capture efficiency and doublet rate of Chigene V2-seq using species mixing. **(C)** Species specificity of unique molecular identifiers (UMIs) in an A549-MC38 cell mixture. Total number of identified A549 cells: n = 2089; total number of identified MC38 cells: n = 1207. The median species specificity of UMIs in A549 cells was 0.954, whereas the median species specificity of UMIs in MC38 cells was 0.932. **(D-E)** Violin plots and box plots depicting the number of UMIs and genes detected in individual A549 and MC38 cells. **(F)** Read distribution along the gene body compared between bulk RNA-seq and three different full-length scRNA-seq methods (Chigene V2-seq, VASA, and snRandom RNA profiling). **(G-H)** Representative raw reads aligned to the human genes GAPDH and ATCB via four different scRNA-seq methods. **(I-J)** Clusters representing subpopulations in A549, 293T, and HeLa cell mixtures in Chigene V2-seq, with the coverage of ATCB c.565G (red) and GAPDH c.463A (blue) detected in each cluster. **(K-L)** Number and percentage of cells with coverage of ATCB c.565G and GAPDH c.463A detected in each cluster. **(M)** Correlations between gene expression measurements via the V2-seq and V1-seq methods. **(N)** UMAPs of single-cell profiles (dots) from human urachal carcinoma libraries colored according to the different platforms used. **(O)** UMAPs showing single-cell profiles (dots) from human urachal carcinoma libraries colored by cell type. **(P)** Proportion of cells of each cell type (y-axis) detected via different methods (x-axis).

The impressive ability of the Chigene V2 platform to detect full-length mRNAs suggests that it may be an ideal method for detecting mutations, gene fusions and alternative splicing at the single-cell level. To improve the sensitivity of detecting mutations, particularly in low-abundance transcripts and rare cells, a nested PCR method was employed for target enrichment of gene mutations. To test this approach, a mixture of 293T, A549, and HeLa cell lines in equal proportions was analysed via Chigene V2-seq to detect SNP sites in two housekeeping genes (ACTB c.565G and GAPDH c.463A). The findings revealed the accurate identification of distinct cell clusters representing the three cell lines, suggesting minimal cross-contamination associated with the Chigene V2 platform (Figure 3I-J). Moreover, the detection of these two SNP sites in the majority of cells underscores the strength of the Chigene V2 platform in conjunction with nested PCR for mutation detection at the single-cell level (Figure 3K-L). Furthermore, by analysing mixtures of non-small cell lung cancer cell lines containing specific mutations (EGFR_p. T790M and KRAS_p. G12S) in varying proportions (Figure S3B-C and supplementary table 2), as well as in a cohort of tumor samples (Figure S3D and supplementary table 3), we found that the detection results closely aligned with the true distributions. Notably, in the Chigene V2 cell line mixing experiments and the Chigene V2 detection of gene mutations in solid tumors, the results indicate that the platform is capable of detecting rare cells, with a detection threshold of approximately 1% or higher (Figure S3E-I). To evaluate whether the Chigene V2 platform could provide sufficient information from clinical tissue samples similar to the Chigene V1 platform, we collected fresh human urachal carcinoma samples and compared their RNA profiles via Chigene V1-seq and Chigene V2-seq. The results revealed a strong correlation between the total RNA profiles obtained via Chigene V1-seq and those obtained via Chigene V2-seq (Spearman R: 0.86, Figure 3M).

Furthermore, the majority of cell types were consistently identified by both methods in the same sample (Figure 3N and Figure S3J-K). Unsupervised clustering analysis identified ten types of cells in the merged sequencing data (Figure 3O and Figure S3L). No significant differences in the proportions of cell types were detected between the V2-seq and V1-seq results (Figure 3P).

### Identification of multi-landscape by Chigene V2 platform in the clinical samples

To assess the performance of the Chigene V2 platform in exploring alternative gene splicing, mutation, and fusion, fresh human urothelial cancer samples were subjected to Chigene V2-seq, whole-exome sequencing (WES) and bulk RNA-seq. A total of 4452 cells were identified from the single-cell suspension, with an average of 1696 genes and 4665 UMIs detected per cell. These cells were clustered into seven distinct cell types (Figure 4A and Figure S4A). By profiling full-length transcripts via the Chigene V2-seq platform, alternative splicing (AS) patterns across different cell types were identified through the quantification of inclusion rates of nonoverlapping exonic regions. In this sample, seven different AS events (total: 3495 events) were detected (Figure 4B), mainly in epithelial cells and T cells (Figure 4C). Notably, alternative first exon (AFE) events were frequently detected (Figure 4B-C), further confirming the high coverage of mRNAs in the Chigene V2 platform. Using the combined WES and RNA-seq data, ten genes with relatively high mutation frequencies at both the DNA and RNA levels were selected for targeted enrichment via the Chigene V2 platform (Figure S4C). UMAP visualization revealed a multitude of gene mutations and comutations in epithelial cells (Figure 4D). To substantiate the reliability of the Chigene V2 platform and investigate the functions of the identified mutated genes at the single- cell level, particular attention was given to the CDKN1A p.F53Ffs*10 mutation (Figure 4E). Compared with those in CDKN1A wild-type epithelial cells, gene ontology (GO) and KEGG enrichment analyses revealed significant enrichment of upregulated genes related to cell cycle- related biological processes (Figure 4F) and signalling pathways (Figure 4G) in CDKN1A- mutated epithelial cells. Cell communication analysis revealed that, compared with wild-type cells, CDKN1A-mutated epithelial cells had greater interactions with endothelial cells (Figure 4H). Specifically, CDKN1A-mutated epithelial cells presented increased VEGF signalling to endothelial cells (Figure 4I), suggesting a potential role in promoting angiogenesis. Similar conclusions supporting the single-cell findings were obtained via TCGA data analysis (Figure S4D-E).

**Figure 4.**
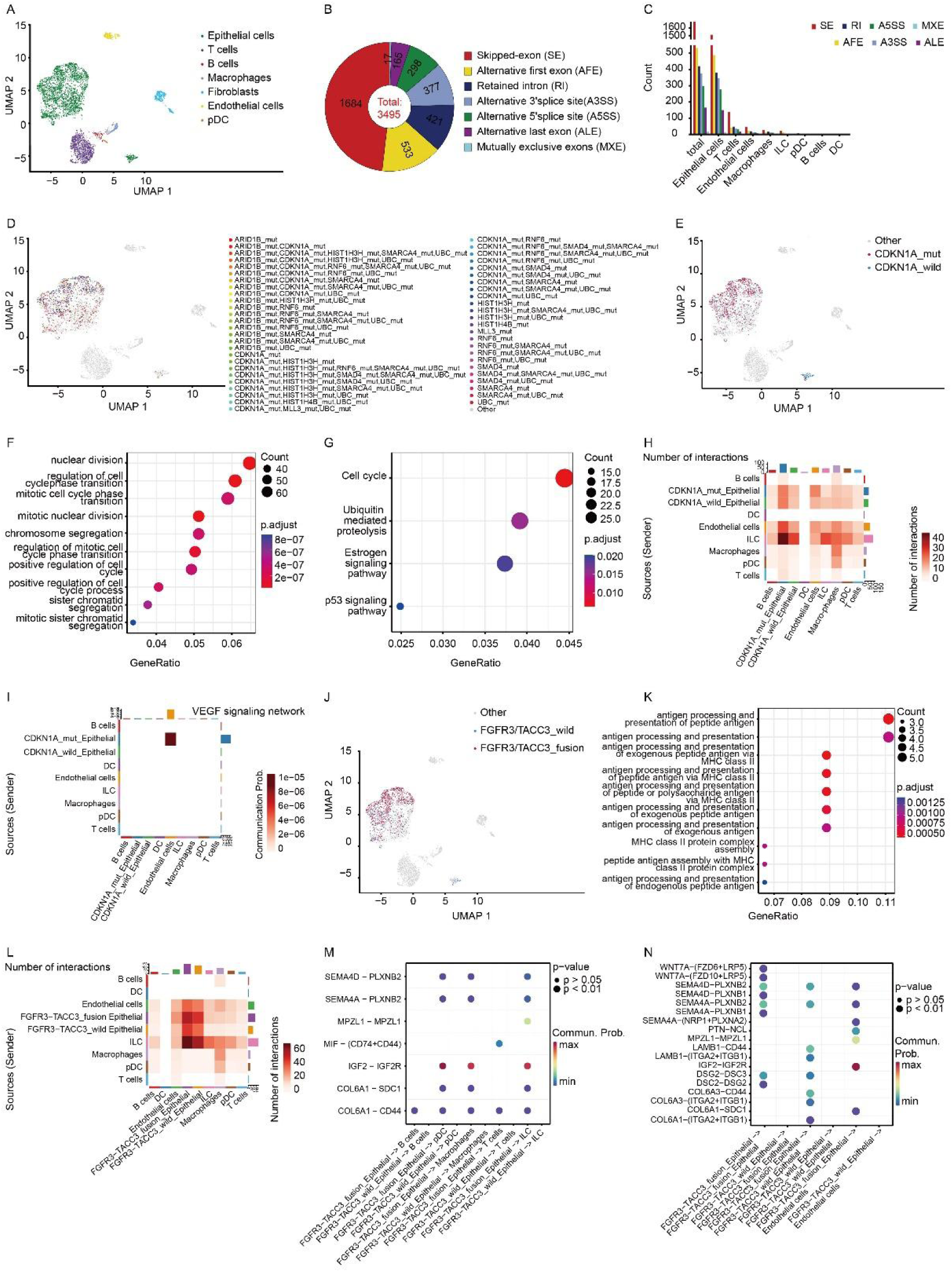
Identification of multi-landscape in clinical samples via the Chigene V2 platform. **(A)** UMAP representation of cells from the human urothelial cancer sample based on their gene expression, with seven annotated cell types displayed. **(B)** Pie plot illustrating the number of detected global gene AS events in Chigene V2-seq. **(C)** Abundance analysis of different classes of AS events across cell clusters. **(D)** Landscape visualization of mutated genes at the single-cell level in human urothelial cancer. **(E)** Detection of CDKN1A mutations in epithelial cells via Chigene V2-seq. **(F-G)** GO and KEGG enrichment analyses of upregulated genes in CDKN1A-mutated cells. **(H)** Heatmap displaying the total number of interactions among different cell populations. **(I)** Heatmap showing the specific VEGF signaling network between CDKN1A-mutated epithelial and endothelial cells. **(J)** Detection of the FGFR3-TACC fusion gene in epithelial cells via Chigene V2-seq. **(K)** GO enrichment analysis of genes downregulated in FGFR3-TACC fusion cells. **(L)** Heatmap revealing the total number of interactions among different cell populations. **(M-N)** Bubble diagrams showing the probability of communication and statistical significance of receptor‒ligand pairs between FGFR3-TACC fusion/wild-type epithelial and immune/nonimmune cells. The dot color represents the communication probability, reflecting the likelihood of successful information exchange between these cell types. The dot size represents the computed p value, which is used to assess the statistical significance of the observed interactions. Empty spaces indicate a communication probability of zero. P values were computed via a one-sided permutation test.

Next, we explored the fusion gene in the Chigene V2 platform and identified the fusion gene FGFR3-TACC in the single-cell dimension (Figure 4J and Figure S4B). GO enrichment analysis revealed that downregulated genes in FGFR3-TACC fusion epithelial cells were significantly enriched in antigen processing and presentation-related biological processes (Figure 4K), potentially indicating a contribution to immune evasion. Cell communication analysis revealed greater signal output from FGFR3-TACC fusion epithelial cells than from wild-type FGFR3 epithelial cells (Figure 4L). Ligand‒receptor pair analysis revealed that FGFR3‒TACC fusion epithelial cells preferentially transmitted signals to both immune cells and nonimmune cells via immunosuppressive ligand‒receptor pairs and promoted angiogenesis and tumor proliferation ligand‒receptor pairs (Figure 4M‒N), such as SEMA4D‒PLXNB1/2^[23–27]^, COL6A1‒SDC1‒CD44^[28]^, MPZL1‒MPZL1^[29–31]^, and WNT7A‒FZD6/10+LRP5) ^[32]^. These findings suggest that the FGFR3-TACC fusion might hinder the efficacy of immunotherapy and contribute to poor prognosis by affecting both immune cells and nonimmune cells. Similar conclusions supporting the single-cell findings were obtained via TCGA data analysis (Figure S4F-G). In summary, the Chigene V2 platform has the ability to explore alternative splicing, mutations, and fusions at the single-cell level, and the results obtained are robust.

### Identification of novel mutated genes via the Chigene V2 platform

To identify novel mutated genes, a fresh sample of urothelial carcinoma was analysed via Chigene V2-seq. A total of 4491 cells with an average of 1821 genes and 4273 UMIs per cell were identified. Unsupervised clustering revealed several distinct clusters representing different cell types, including DCs, endothelial cells, epithelial cells, T cells, ILCs, and macrophages (Figure 5A and Figure S5A). Analysis of the Chigene V2-seq data revealed multiple gene mutations in epithelial cells (Figure 5B). Among these, we specifically focused on the ARHGAP5 gene, which harbors the p.I307V mutation, owing to its high mutation frequency and the unclear biological function and clinical significance associated with this alteration (Figure 5C). Sanger sequencing validated the true existence of the mutation site in this gene (Figure S5B). GO enrichment and alternative splicing analyses demonstrated a greater frequency of RNA splicing events in ARHGAP5-mutated epithelial cells (Figure 5D-E). Cell communication analysis further revealed that epithelial cells with mutated ARHGAP5 displayed a distinct number of interactions with other clusters, particularly notable in their interaction with T cells (Figure 5F-G). Subsequent examination of ligand ‒ receptor pairs revealed notable differences between ARHGAP5-mutated and wild-type tumor cells regarding the signals they transmitted to both immune and nonimmune cells (Figure 5H). In contrast to ARHGAP5 wild-type epithelial cells, ARHGAP5-mutated epithelial cells tend to transmit signals to dendritic cells (DCs) via the F11R‒F11R interaction, which can impede DC migration to lymph nodes and the initiation of tumor-specific immune responses ^[33]^. Additionally, in ARHGAP5-mutated epithelial cells, signals are specifically transmitted to T cells through APP-CD74, which is known to exert an inhibitory effect on the activation of CD4+ T cells (Figure 5I). Moreover, ARHGAP5-mutated epithelial cells also transmit signals to nonimmune cells involved in tumorigenesis, metastasis, and angiogenesis (Figure 5J) via MPZL1-MPZL1^[29–31]^, F11R-F11R^[34]^, and COL6A1-SDC1^[28]^. Consequently, it is speculated that urothelial carcinoma patients with ARHGAP5 mutations may exhibit a poorer response to immunotherapy. Additionally, TCGA data analysis (Figure S5C) and validation through cell lines (Figure S5D) supported the findings related to the ARHGAP5 mutation. Overall, our experimental results corroborate the findings from the TCGA analysis and provide further validation of the role of the ARHGAP5 p.I307V mutation in promoting pathways that enhance tumor malignancy, metastatic potential and immune evasion. Furthermore, these findings validate the functionality of our Chigene V2 platform, demonstrating its ability to provide insights into the functional implications of this mutation at the single-cell level, underscoring its importance in tumor biology.

**Figure 5.**
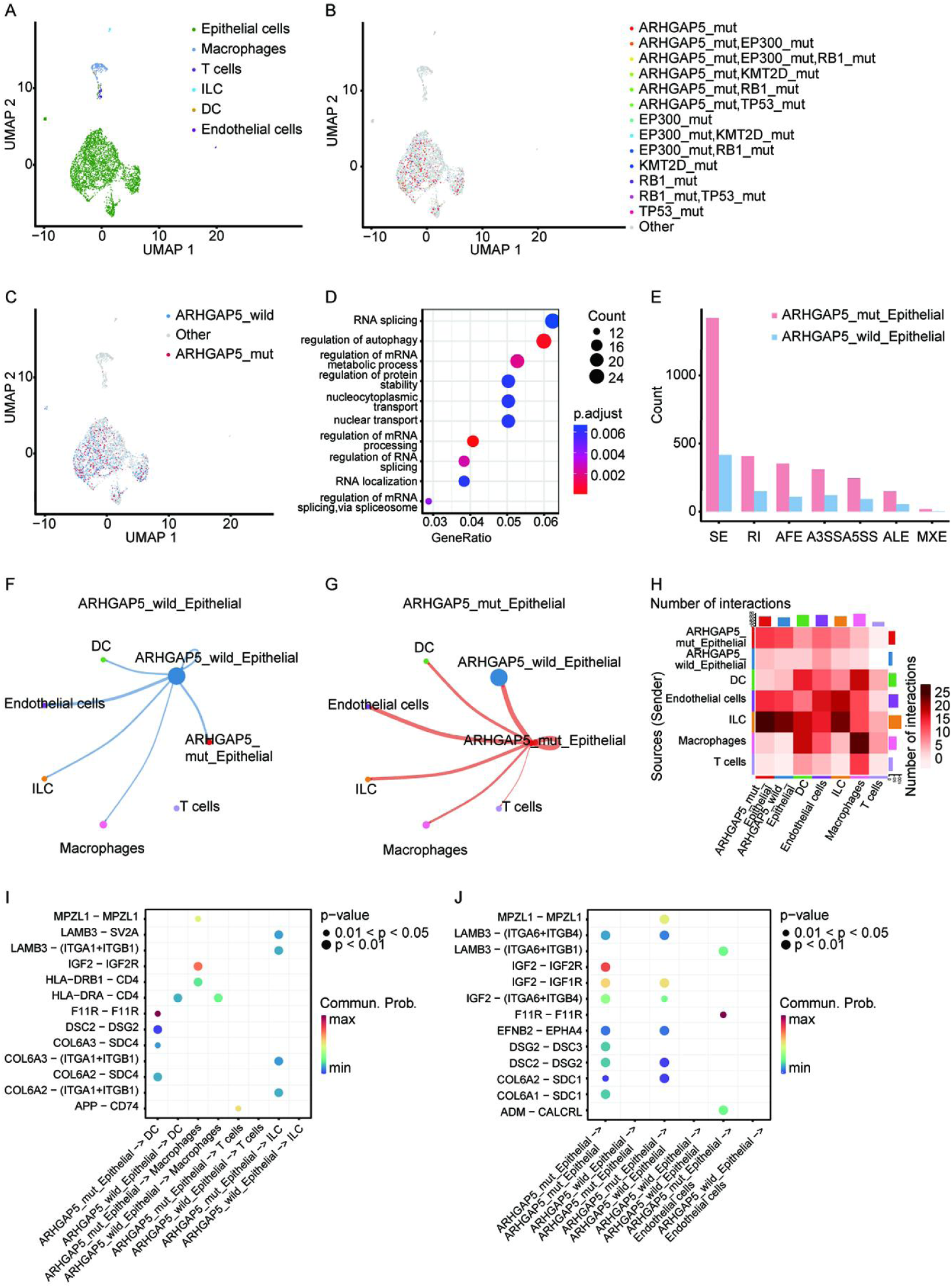
Chigene V2 enables the discovery of novel gene mutations in urothelial carcinoma at the single-cell level. **(A)** UMAP visualization of genes expressed in cells from a human urothelial cancer sample, with six annotated cell types displayed. **(B)** Landscape depiction of mutated genes at the single-cell level in human urothelial cancer. **(C)** Detection of the ARHGAP5 mutation in epithelial cells via Chigene V2-seq. **(D)** GO enrichment analysis of upregulated genes in ARHGAP5-mutated cells. **(E)** Comparison of AS events between ARHGAP5-mutated and wild-type cells. **(F-G)** Total number of interactions between ARHGAP5-mutated/wild-type cells and other cell populations. The circle sizes represent the number of cells in each cell group, whereas the edge width represents the communication probability. **(H)** Heatmap displaying the total number of interactions among different cell populations. **(I-J)** Bubble diagrams showing the communication probability and statistical significance of receptor‒ligand pairs between ARHGAP5-mutated/wild-type epithelial cells and immune/nonimmune cells. The dot color represents the communication probability, reflecting the likelihood of successful information exchange between these cell types. The dot size represents the computed p value, which is used to assess the statistical significance of the observed interactions. Empty spaces indicate a communication probability of zero. P values were computed via a one-sided permutation test.

To gain a more comprehensive understanding of the prevalence of ARHGAP5 mutations across various cancers, we searched the TCGA database (Figure S5E). Intriguingly, our analysis revealed a mutation frequency of approximately 2.6% for ARHGAP5 in urothelial carcinoma (Figure S5F). To assess the clinical implications of these mutations, propensity score matching (1:1) was conducted, matching patients with ARHGAP5 mutations to their wild-type counterparts based on variables such as cancer type, age, sex, race, and pathological stage (Figure S5G). Despite not reaching statistical significance, an intriguing pattern emerged: progression-free survival (PFS) tended to be shorter in the mutant patients (Figure S5H). The associations between ARHGAP5 mutations and immunotherapy response in cohorts of patients who underwent immunotherapy for urothelial carcinoma^[35]^ or advanced clear cell renal cell carcinoma^[36]^ were further investigated. Interestingly, no ARHGAP5 mutations were detected among the 9 responders in the urothelial cancer cohort, whereas one such mutation was present among the 16 nonresponders. In the advanced clear cell renal cell carcinoma cohort, ARHGAP5 mutations were identified in 5 out of 261 patients. Among these patients, 2 had stable disease (SD), and 3 experienced progressive disease (PD). The distribution of these clinical responses indicates a possible association between ARHGAP5 mutations and unfavourable treatment outcomes. By integrating data from the TCGA database, immunotherapy cohorts, and WES analysis, we propose that ARHGAP5 gene mutations might be linked to decreased progression-free survival (PFS) and poor response to immunotherapy.

## Discussion

In this study, we introduced a microfluidic chip-based high-throughput single-cell RNA sequencing (scRNA-seq) system designed for comprehensive analysis of mRNA expression, mutations, alternative splicing, and even gene fusions at the single-cell level. Our scRNA-seq platform consists of two versions. Compared with mainstream platforms such as 10X Genomics, BD Rhapsody, and Singleron, Chigene V1, a conventional single-cell sequencing platform utilizing 3’ end sequencing, has demonstrated equivalent performance and scalability. Chigene V2 specifically captures mRNAs with more uniform coverage across the gene body and achieves greater sequencing depth for coding sequences. Notably, our Chigene scRNA-seq platform employs standard molecular biology procedures and a well-established microwell barcoding platform. As a result, the Chigene scRNA-seq platform is user friendly and can be readily used for large-scale applications.

Most contemporary single-cell RNA sequencing (scRNA-seq) platforms depend primarily on the hybridization of barcoded oligo-dT primers. However, this approach limits the ability of these methods to detect short fragments (∼400--600 base pairs) near the poly(A) tail, thereby hampering the thorough investigation of gene mutations, gene fusion, and alternative splicing at the single-cell level. Recent advancements in total RNA full-length scRNA-seq methods have addressed these challenges ^[18, 20, 37]^. However, the use of total RNA sequencing techniques presents inherent limitations in sequencing depth dedicated to mRNAs owing to the presence of noncoding RNAs and alternative splicing variants in the total RNA pool. Consequently, these molecules can compete for sequencing data, resulting in a reduced sequencing depth for mRNA detection. The Chigene V2 platform introduces a novel random bridging colabelling technique, facilitating the specific capture of full-length mRNAs. This method demonstrated improved alignment with conventional 3’ single-cell sequencing, effectively minimizing interference from rRNA and other transcripts. This innovation eliminates the need for additional bioinformatics or enrichment methods to remove rRNA, thereby saving time and experimental costs. Furthermore, data utilization can be enhanced by increasing the sequencing depth for mRNAs within the same data volume. In addition, by incorporating targeted capture techniques, our approach facilitates the detection of low-abundance mutations and fusions, which substantially enhances sensitivity (Figure S6A-B). This combination of specific mRNA capture and targeted mutation detection results in notable improvements in both the breadth and sensitivity of our platform, allowing for a more comprehensive analysis of mutations and fusions within the transcriptome. These innovations underscore the novelty and effectiveness of our methods in advancing transcriptome-level research.

The theoretical throughput of Chigene V2 is equivalent to that of conventional 3’ single-cell sequencing. Among the existing single-cell sequencing platforms, the Chigene V2 platform is superior for mRNA analysis because of its more even gene body coverage and sequencing depth. These advantages enhance the analysis of alternative splicing in single-cell polyadenylated RNA species and facilitate the identification of rare subpopulations for precision diagnosis and treatment of human diseases. Moreover, when combined with targeted capture, the Chigene V2 platform achieves increased sensitivity for detecting gene mutations and gene fusions. These capabilities enable the simultaneous exploration of gene expression, gene mutations, alternative splicing, and gene fusion at the single-cell level.

To validate the potential of our platform for multidimensional clinical sample testing, we utilized our Chigene V2 platform to concurrently detect mutations, alternative splicing, and gene fusions in several human cancer samples. When we investigated known functional gene mutations, such as CDKN1A, and gene fusions, such as FGFR3-TACC, via the Chigene V2 platform, the results implied that enrichment analyses of differential gene expression between wild-type and mutant cells were consistent with previously reported functional changes in tumor cells. Additionally, our analysis revealed that intercellular signalling can affect the TME, which deepened our understanding of genes with known functions. Notably, in this study, we identified a novel gene mutation site in ARHGAP5, which has not been previously reported in the context of tumors. We utilized this example to illustrate the finer resolution of our Chigene V2 platform in bladder cancer research. Importantly, through TCGA database analyses and experimental validation via cell line studies, we further demonstrated that ARHGAP5 functions as a tumor suppressor gene, potentially regulating tumor cell malignancy and immune evasion. Additionally, analyses from the TCGA database indicate that this mutation has potential as a prognostic biomarker across various cancers (Figure S5). Interestingly, more recently, ARHGAP5 has also been implicated as a tumor suppressor in colorectal cancer, where its mutation is associated with shortened progression-free survival (PFS)^[38]^. These findings reinforce the potential of our single-cell findings to reveal novel gene mutations and their roles in cancer. We plan to conduct more detailed mechanistic studies and animal model validations to assess the impact of the ARHGAP5 mutation on bladder cancer comprehensively.

Nevertheless, it is important to acknowledge certain limitations of our technique. First, like other full-length transcriptome sequencing technologies, Chigene V2 currently supports the detection of mutations only in expressed genes. Mutations present in non-expressed genes currently remain undetectable. However, it is worth reconsidering whether these nonexpressed mutations may nonetheless influence tumor development and progression, as this limitation may not have a profound impact on the overall applicability of our platform. Second, the single-cell cDNA amplification method relies on poly(T) primers, which restrict the detection of mRNA molecules lacking a poly(A) tail, such as histone mRNAs^[39]^. Consequently, these specific transcripts were not captured or detected by our mRNA-seq assay. Moreover, given that our technology captures a broader range of mRNA regions, the data volume required for analysis is relatively greater than that of some existing platforms. This aspect should be considered when evaluating the practicality of implementing Chigene V2 in various research settings. Finally, as our assay utilizes mature mRNAs as templates, mutations present within intronic and intergenic regions are not detected. These limitations could be addressed through future technical refinements.

In conclusion, the straightforward experimental protocols and comprehensive transcriptomic information offered by our platform show promising potential for the significant contributions of Chigene scRNA-seq to large-scale applications in both basic and clinical research in the future. The promotion and application of our platform are expected to advance the precision of personalized medicine and therapeutic target discovery by providing deeper insights into the genetic landscape of tumors.

## Method

### Microfluidic chip device fabrication and microbeads

The chip comprises a support layer, a chip microstructure layer, and a chip top cover layer. The support layer was composed of rigid glass, whereas all other layers were constructed using polydimethylsiloxane (PDMS) material with mold casting. For the chip microstructure layer, a photomask was created based on the chip design, and a mold was fabricated via traditional photolithography methods (with an SU-8 negative photoresist). The resulting mold consisted of an array of cylindrical microstructures with dimensions of 52 μm (height) and 38 μm (diameter) and a center-to-center distance of 48 μm between cylinders. The PDMS mixture, consisting of a PDMS base and curing agent at a ratio of 10:1, was adequately mixed and poured onto the chip microstructure layer mold. After baking at 80°C for 30 min, the cured PDMS chip microstructure layer was gently removed from the mold. Moving to the chip top cover layer, a mold was prepared on the basis of the chip design, including a chip chamber framework section and a top cover section. Similarly, PDMS was cast onto the mold to form the chip top cover layer. To complete the chip assembly, 1 mm diameter holes were punched in the top cover section, which served as the inlet and outlet ports. The glass layer, PDMS chip microstructure layer, and PDMS chip top cover layer were subsequently treated with plasma and bonded together to form the final microwell. The microbeads were purchased from Chemgenes Corporation (MACOSKO-2011-10 B) and were filtered through a 40 µm strainer. A detailed protocol for the Chigene V1 assay, including the oligo list, is provided in the Supplementary Materials (Supplementary Methods and Supplementary Table 1).

### Species mixture experiment and human samples

HeLa cells, A549 cells, 293T cells, and MC38 cells were obtained from the CAMS Cell Culture Center and cultured in Dulbecco’s modified Eagle’s medium (DMEM, Gibco, Cat #11965092) supplemented with 10% heat-inactivated fetal bovine serum (FBS, Gibco, Cat # 26010074). The cells were maintained at 37 °C in a 5% CO2 incubator (Thermo Heracell 240i) and passaged every 2 days. For the species mixture experiment, A549 and MC38 cells were harvested and washed three times in PBS by centrifugation at 4 °C and 600 × g for 3 min. Subsequently, the cells were individually counted, and equal numbers of cells from each cell line were mixed. The resulting cell mixture was subjected to single-cell RNA sequencing (scRNA-seq) following the Chigene scRNA-seq protocol. The collection of human peripheral blood and urological cancer samples, as well as the research conducted in this study, was approved by the Research Ethics Committee of Sun Yat-sen Memorial Hospital, Sun Yat-sen University (approval numbers: SYSKY-2023-396-01). Clinical information was collected after written informed consent was obtained.

### Library preparation

The chip was first preprocessed using PBS I buffer (PBS + RNase inhibitor), and then approximately 15,000 cells and capture beads were loaded onto the chip in turn. After the redundant cells and beads were washed out, cell lysis occurred in the microwells of the chip. The RNA molecules released from the lysed cells were captured by beads coated with oligo sequences in the same microwell. Then, the lysis buffer in the chip was rapidly exchanged with air and high-density wash buffer, and all the beads, including the mRNA-captured beads, were recovered by washing. Approximately 80% of the beads were collected by two washes, and additional washing steps helped to obtain more beads. Next, cDNA molecules with cell barcodes and UMI sequences were obtained through reverse transcription. After ExoI digestion and amplification, the full- length cDNA was enriched, and a single-cell RNA sequencing (scRNA-seq) library was constructed via modified Tn5 transposase-based methods. Briefly, 5–10 ng of cDNA was fragmented and amplified to construct the sequencing library. This approach will help to obtain more information on the gene body for mRNA isoform analysis. Sequencing was performed via PE150 bp on the MGI-T7 platform (MGI, China).

### Random bridging colabelling for Chigene V2-seq

To obtain full-length mRNA, after reverse transcription and template switching, the mRNA- free barcode oligo was annealed with the tag primer via the following reaction mixture: 100,000 microbeads in 35 μl of NF-water, 5 μl of 10 μM tag primer, and 10 μl of 20X SSC Buffer (NEB).

The reaction mixture was incubated at room temperature for 15 minutes. After annealing, the microbeads were washed with PBS. To modify the 3’ end of the barcode oligo, an extension mixture consisting of 2 μl of NEB Buffer 10X, 5 μl of dNTPs, 2 μl of Klenow (exo-), and 11 μl of NF-water was added. The mixture was then incubated at 37°C for 30 minutes. Denaturing buffer (Illumina) was added to remove the mRNA and tag primer from the microbeads. The microbeads were then washed twice with PBS. Bridge oligo1 and bridge oligo2 were annealed to form a bridge dimer. One end of the bridge dimer was reverse complemented to the modified barcode oligo, whereas the other end contained tens of random nucleotides "N" at the 3’ tail. When the modified barcode oligo annealed with the bridge dimer, the 3’ end of the barcode oligo joins the neighboring 5’ end of the N-tail hand. The bridge capture mixture, which consisted of 10 μl of 20X SSC buffer, 10 μl of 10 μM bridge dimer, and 100,000 microbeads in 30 μl of NF-water, was incubated at room temperature for 15 minutes. The microbeads were then washed with PBS and suspended in the ligation reaction mixture, which included 2 μl of 10X T4 DNA ligase buffer, 5 μl of ATP, 2 μl of T4 DNA ligase, and 11 μl of NF-water. This mixture was incubated at room temperature for 30 minutes. After washing with PBS, extension was performed via the following reaction mixture: 2 μl of NEB Buffer 10X, 5 μl of dNTPs, 2 μl of Klenow (exo-), and 11 μl of NF-water. To improve efficiency, repeating the steps from bridge capture to extension 2--3 times is recommended. To block the free barcode oligo, ddNTPs were used, and then the cDNA was amplified and purified.

### Targeted capture of specific regions of genes

For the confirmation of mutation sites or gene fusions by WES, nested PCR primers were designed upstream and downstream of each target region. The upstream-out primers, downstream- out primers, upstream-in primers, and downstream-in primers were mixed separately. A total of 2.5 ng of cDNA was used as a template for amplification with the upstream and downstream primers. Subsequently, secondary PCRs were carried out using the upstream and downstream primers. After each round of amplification, the reaction products were individually purified via magnetic beads. The purified products from each reaction were then combined. Index PCR was performed on the combined products to generate a sequencing library. All the libraries were subjected to quality control analysis via an Equalbit 1× dsDNA HS Assay Kit (Vazyme, China) and a Standard (S2) Cartridge Kit (Bioptic, Taiwan, China). This quality control ensures that the libraries meet the desired standards for further downstream analysis.

## Data analysis

### Preprocessing of the Chigene V1/V2-seq data

First, the raw read data were preprocessed via a Python script. For each Read1, the cell-specific barcode (16 nucleotides) located at the beginning of the read was identified. Hamming distance correction was applied by comparison to a whitelist of 100,000 barcodes provided by the Chigene platform. Barcodes within the whitelist or with a Hamming distance of 1 or less were considered valid cell barcodes and retained for downstream analysis. Barcodes not present in the whitelist or with a Hamming distance exceeding 1 were discarded to ensure accurate cell identification. The corrected barcode from Read1, along with the UMI (8 nucleotides), was appended to the read ID of Read2. The resulting reads were aligned to the reference genome (Chigene V1: GRCh38, Chigene V2: GRCh37, mixed human‒mouse genome: GRCh38 and the GRCm39 hybrid genome) via the STAR aligner (version 2.7.1a) with default parameters. Reads uniquely mapped to the reference genome were assigned to genes via featureCount (version 2.0.1). UMItools (version 1.0.1) was used to correct UMI errors and perform UMI quantification within individual cells, resulting in the generation of the raw matrix. The EmptyDrops cell calling algorithm was utilized to identify true cells from the background and generate a gene expression matrix for subsequent analysis.

### Clustering and downstream analysis

The single-cell expression matrix was imported into Seurat (version 3.0.2). Cells with fewer than 500 detected genes or more than 20% mitochondrial UMI counts were filtered out. Genes detected in more than 3 cells were retained. The genes were then ranked in descending order based on residual variance estimated via the "vst" method implemented in the FindVariableFeatures function of Seurat. The top 2000 genes, identified as highly variable genes (HVGs), were selected for further analysis. Principal component analysis (PCA) was performed on these highly variable genes. A shared nearest neighbor (SNN) graph was constructed using the top 15 principal components, and the cells were clustered via the Leiden algorithm. The resolution parameter for clustering was set to 0.6, while all other parameters were kept at the default values. To visualize the clusters, uniform manifold approximation and projection (UMAP) was applied in Seurat. After completing the clustering analysis, the cell types were annotated via a cell type (version 0.1.9).

### Expression correlation analysis for the Chigene V1/V2 platform

To compare the gene expression correlation between the Chigene V1 and Chigene V2 platforms, we imported the expression matrices of both platforms into Seurat (version 3.0.2) and performed data normalization and standardization using the same default parameters. We then utilized the Seurat function “AverageExpression” to calculate the average expression levels of genes in each platform. Finally, we used the “cor” function in R (version 4.0.2) to calculate the Spearman correlation coefficient based on the natural logarithm of the average expression levels.

### Read coverage analysis of the gene body

The public VASA-seq data can be accessed at the Gene Expression Omnibus (GEO) under the accession code GSE176588. The public snRandom-seq data can be obtained from the Genome Sequence Archive (GSA) with the accession code "HRA003712". The 10x data were obtained from the PBMC data sequenced by Chigene via the 10x platform. The bulk RNA-seq data were obtained from transcriptome sequencing data. All the data sets were aligned to the GRCh38 reference genome via the STAR Aligner (version 2.7.1a) with the same parameters, resulting in BAM files. For all single cells, the BAM files were used as bulk data. RSeQC (version 4.0.1) was used to calculate the read coverage over the gene body.

### Variant calling

The barcode corrected from Read1, along with the UMI (8 nucleotides), was added to the read ID of Read2. These reads were aligned to the GRCh37 reference genome via STAR (version 2.7.1a) with default parameters. To remove duplicates, the umi_tools’ group and dedup functionalities were applied based on genomic coordinates and UMIs. This process helps eliminate PCR amplification biases and improves the accuracy of variant detection. For variant calling, BCFtools (version 1.9) was utilized to call variants at the single-cell level. The identified mutations were subsequently annotated via ANNOVAR to obtain additional information regarding mutation characteristics and potential functional impacts. Cells that harboured detected mutations were classified as mutant cells, whereas cells without detected mutations were defined as wild-type cells.

### Cell interaction analysis

Cell communication analysis was performed via the R package CellChat (version 1.5.0). After preprocessing the single-cell RNA sequencing data by applying quality control measures and normalizing expression values, we utilized CellChat to identify ligand‒receptor pairs between different cell populations within the TME. The default parameters provided by the CellChat package were employed for comprehensive coverage of potential cell communication events. The probabilities of communication between cell types were estimated based on the identified ligand‒receptor pairs, which represent the likelihood of effective information exchange. Signalling pathways involved in intercellular communication were inferred via the built-in algorithms provided by CellChat. This comprehensive analysis aimed to elucidate the intricate landscape of cell‒cell interactions within the tumor microenvironment and shed light on potential communication between different cell populations.

### GO and KEGG analyses

For the enrichment analysis of DEGs between comparison groups, the FindMarkers function in Seurat was used. This function allows the identification of differential gene sets based on specified criteria such as the percentage of cells expressing the gene (pct) and the p value (p_val). For example, genes with pct > 0.1 and p_val < 0.05 were selected. To perform Gene Ontology (GO) and Kyoto Encyclopedia of Genes and Genomes (KEGG) enrichment analyses of the DEGs, the R package clusterProfiler was used. This package provides functions for functional enrichment analysis and visualization. By default, the package uses hypergeometric tests for enrichment analysis and includes a set of predefined gene sets from databases such as GO and KEGG.

### Statistical Analysis

Statistical details for each experiment are provided in the figure legends. Pre-processing of the Chigene V1/V2-seq data was described in the Data Analysis section. The single-cell isolation experiment and doublet rate experiment were repeated more than three times independently with similar results. The p value (p) for the Sperman’ s correlation coefficient (R) was computed from a two-sided permutation test. The ggpubr package (0.4.0) in R was used for statistical analysis.

## Supporting Information

Supporting Information is available from the Wiley Online Library or from the author.

## Acknowledgements

This study was supported by the National Key Research and Development Program of China (Grant No. 2022YFC2408300), the National Natural Science Foundation of China (Grant No. 82322056, 82341018, 82072827, 81825016, and 82273421), the Guangdong Basic and Applied Basic Research Foundation (Grant No. 2021B1515020009), the Science and Technology Program of Guangzhou (Grant No. 2023A03J0718, 2024B03J1234, and 2024A04J6558), Fundamental Research Funds for the Central Universities, Sun Yat-sen University (23ykbj002 for Xu Chen), the Guangdong Provincial Clinical Research Centre for Urological Diseases (2020B1111170006), and the Guangdong Science and Technology Department (2020B12060018, 2018B030317001, and 2017B030314026).

## Authors’ contributions

Weiyue Gu, Tianxin Lin, Xu Chen and Ligang Wang conceptualized the study. Ligang Wang, Xiao Wang, and Fangxiao Wang were responsible for the experimental design and execution. Tonglei Guo, Fei Xie, and Xiuxin Li conducted the bioinformatics analysis. Wenlong Zhong, Daqin Wu, and Xu Chen contributed to the clinical tissue selection and PBMC acquisition. Chengjunyu Zhang performed various cellular experiments, including ARHGAP5 gene knockdown and overexpression studies, to further investigate the functional role of ARHGAP5 in cancer biology. Lihua Zhao and Minghui Li performed the data analysis. Lihua Zhao contributed to the overall conceptualization and design of the manuscript and drafted the initial version. Lihua Zhao, Wenlong Zhong and Xu Chen participated in manuscript revisions and discussions related to clinical aspects. Weiyue Gu, Xu Chen and Tianxin Lin provided supervision throughout the project. All the authors have read and approved the final version of the manuscript.

## Data availability

The data that support the findings of this study are available from the corresponding author upon reasonable request.

