## Supplementary Figures for "Identification of Multi-landscape and Cell Interactions in the Tumor Microenvironment through High-Coverage Single-Cell Sequencing"

\# To whom contributed this work equally

* To whom correspondence should be addressed. Weiyue Gu,. Tianxin Lin,. Xu Chen,.

**Supplemental Figure**

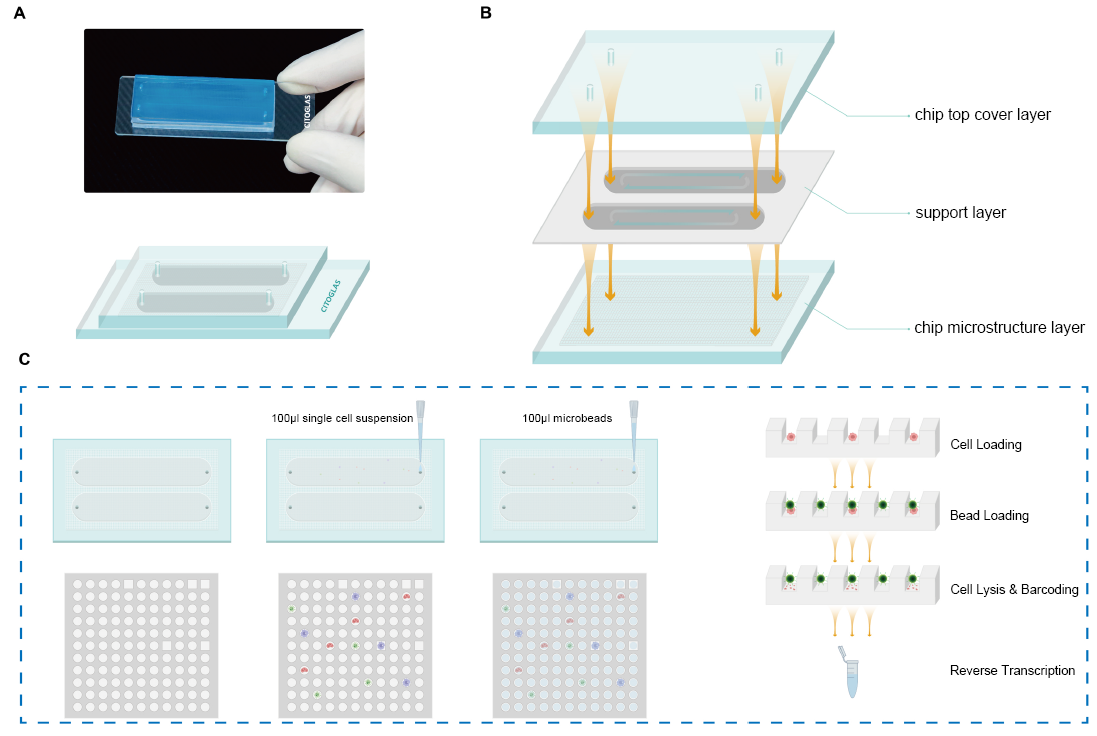

**Figure S1. Overview of the chigene V1 microfluidic chip and library construction process. (A)** Image of the Chigene V1 microfluidic chip, showcasing its design and structure. **(B)** Schematic representation of the chip layers, including the chip top cover layer, support layer, and chip microstructure layer, illustrating how fluid flow facilitates cell distribution. **(C)** Workflow of the library preparation process, detailing the steps of cell loading, microbead loading, cell lysis and reverse transcription, with examples of single-cell suspension and microbead distribution within the micro-wells.

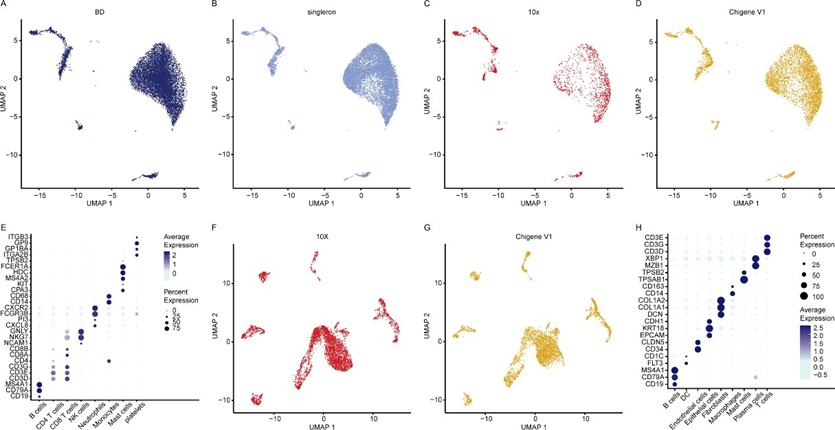

**Figure S2. Comparison of the performances of Chigene V1 and other platforms. (A-D)** Split

UMAP plot of PBMC samples showing consistency between Chigene V1 and other scRNA-seq

platforms. **(E)** Bubble plot showing marker genes of different subpopulations of PBMCs. **(F-G)** Split UMAP plot of urachal carcinoma specimens showing consistency between Chigene V1 and the 10x Genomics platform. **(H)** Bubble plot depicting the expression of marker genes in distinct subpopulations of urachal carcinoma specimens.

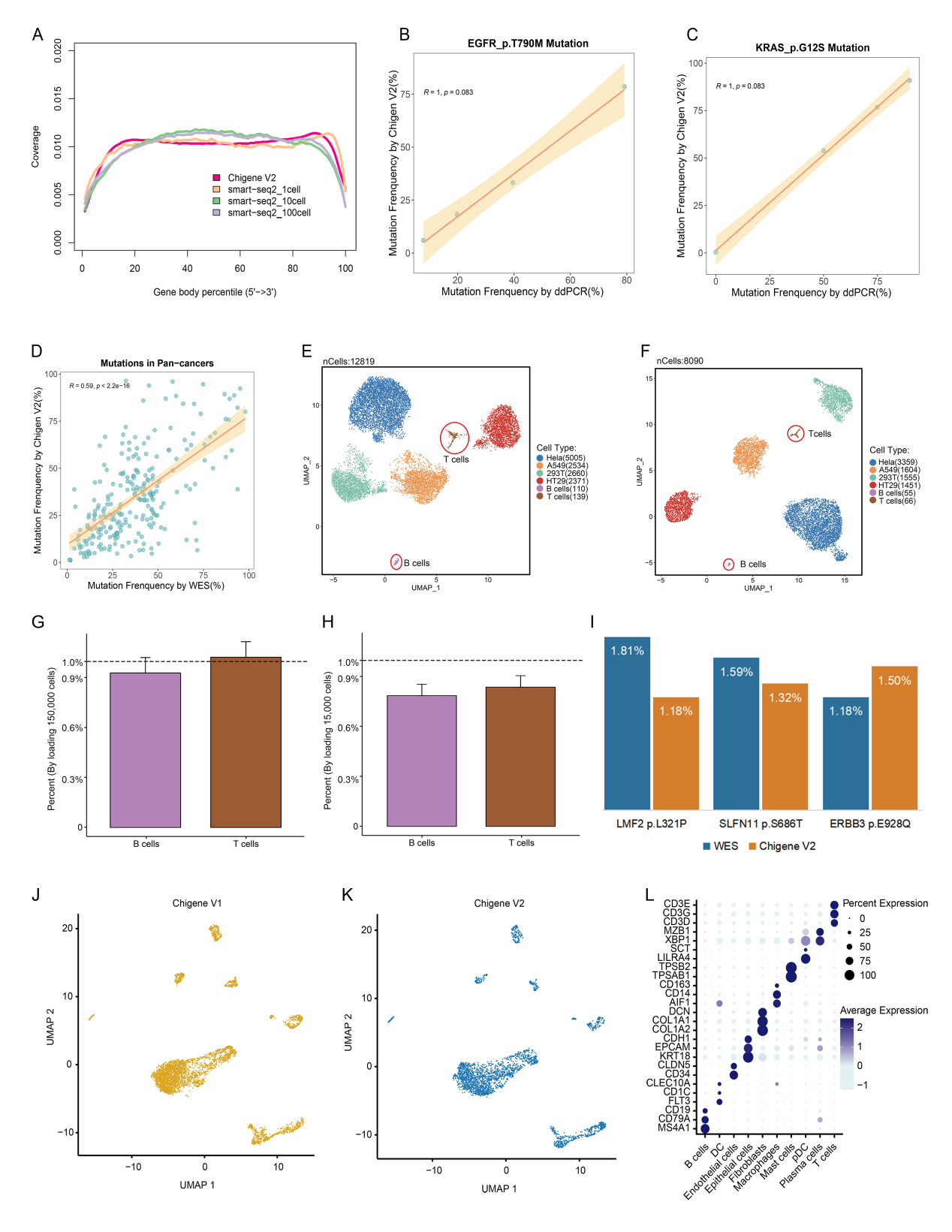

**Figure S3. Development and validation of the Chigene V2 platform. (A)** Coverage uniformity comparison of Chigene V2 and Smart-seq2 across various single-cell conditions. **(B-C)** Comparison of mutation frequency detection for EGFR p.T790M and KRAS p.G12S mutations using the Chigene V2 platform and ddPCR. **(D)** Comparison of mutation detection between Chigene V2-seq and Whole Exome Sequencing (WES) a from various solid tumors. **(E-F)** Split UMAP plot of urachal carcinoma specimens showing consistency between Chigene V1 and Chigene V2. **(G)** Bubble plot depicting the expression of marker genes in distinct subpopulations of urachal carcinoma specimens.

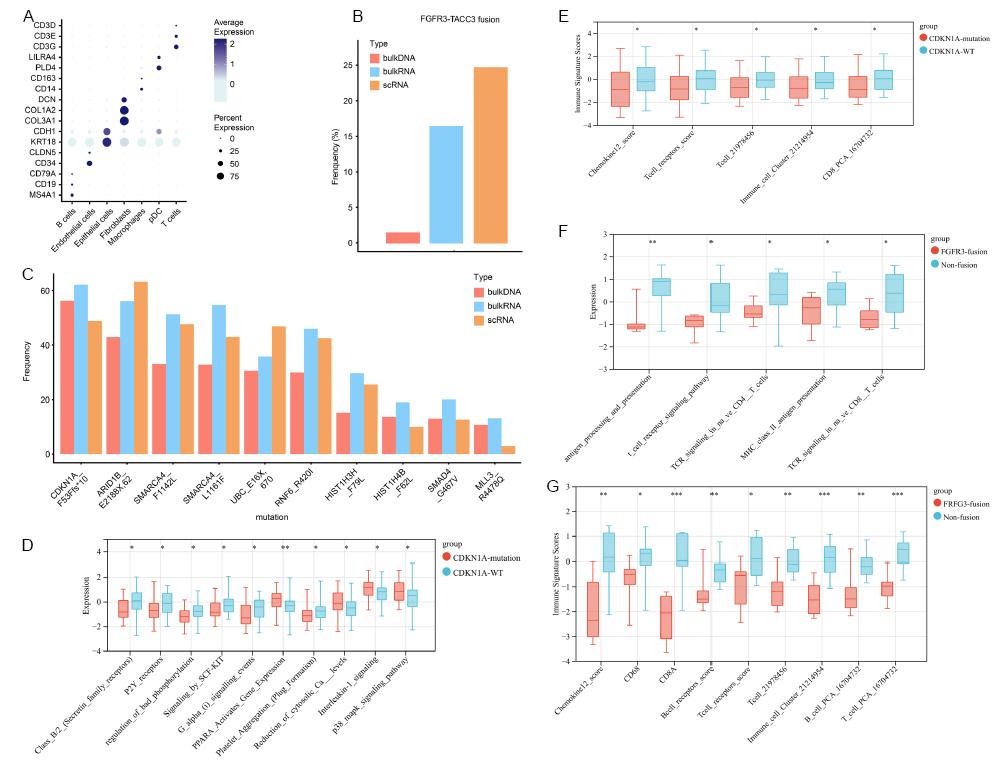

**Figure S4. Identification of multi-landscape in clinical samples via the Chigene V2 platform. (A)** Bubble plot depicting the expression of marker genes in distinct subpopulations of urothelial cancer specimens. **(B)** Frequencies of the FGFR3-TACC3 fusion at the DNA, RNA, and single-cell levels. **(C)** Comparison of mutation frequencies of the top 10 mutated genes identified by whole-exome sequencing (WES) at the DNA, RNA, and single-cell levels. **(D)** Comparison of signaling pathway expression between CDKN1A mutation and wild-type groups in urothelial carcinoma. **(E)** Immune signature scores associated with CDKN1A mutations versus wild-type. **(F)** Expression levels of signaling pathways related to antigen processing and immune response in FGFR3-TACC fusion versus non-fusion groups. **(G)** Immune signature scores in FGFR3-TACC fusion versus non-fusion groups.

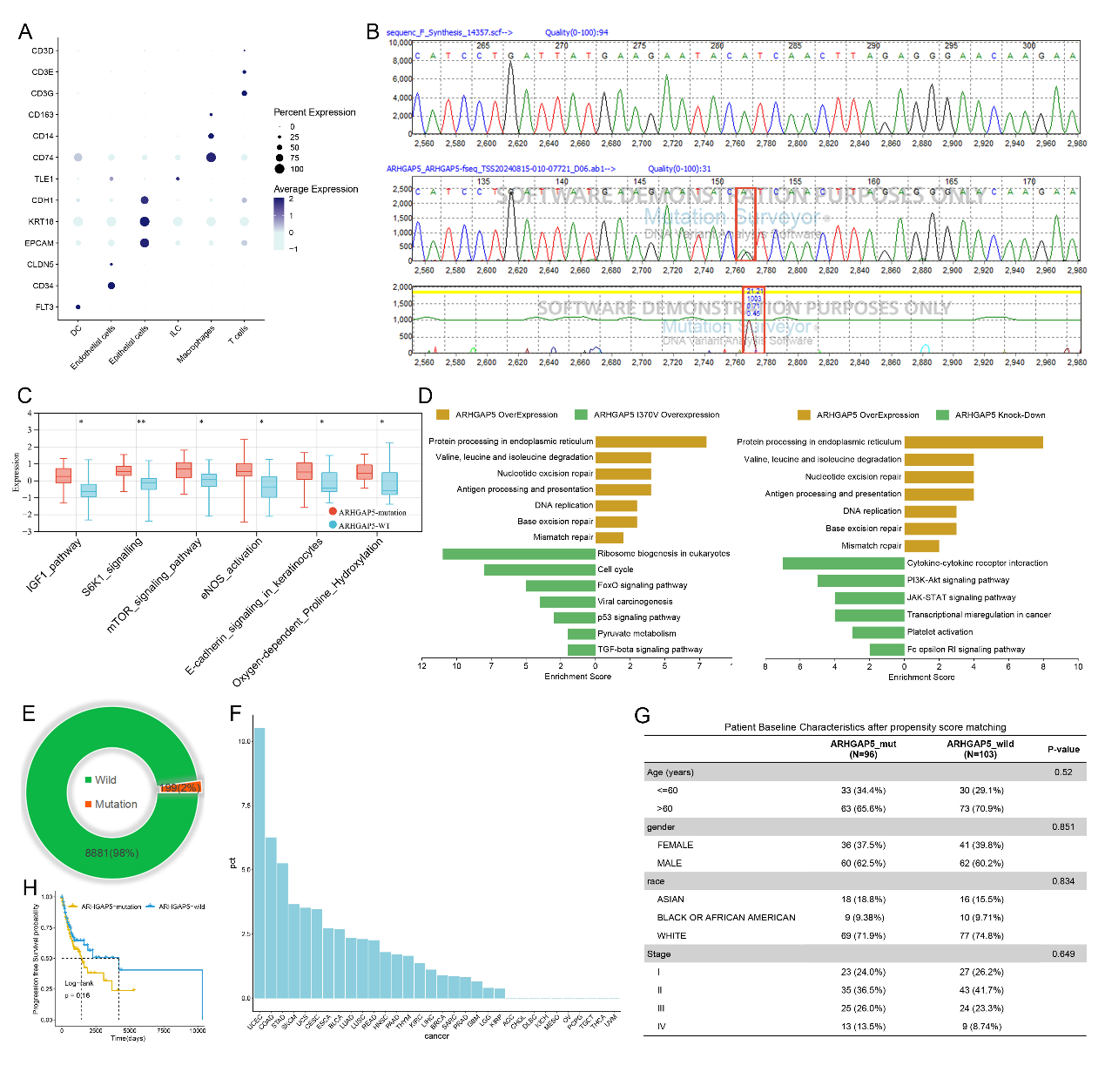

**Figure S5. Identification of novel mutated genes via the Chigene V2 platform.** (A) Bubble plot depicting the expression of marker genes in distinct subpopulations of urothelial carcinoma. (B) Analysis of the ARHGAP5 p.I370V mutation using Sanger sequencing software. (C) Analysis of signaling pathways associated with ARHGAP5 mutations in bladder cancer, derived from the TCGA database. **(D)** Pathway enrichment analysis of ARHGAP5 p.I307V mutation effects from overexpression and knockdown experiments. (E) mutation frequency of ARHGAP5 in Pan-cancer. (F) The mutation frequency of ARHGAP5 in different cancers. (G) Matching propensity scores between ARHGAP5 mutant/wild-type populations. (H) PFS comparison between the ARHGAP5 mutant and ARHGAP5 wild-type groups in Pan-cancer.

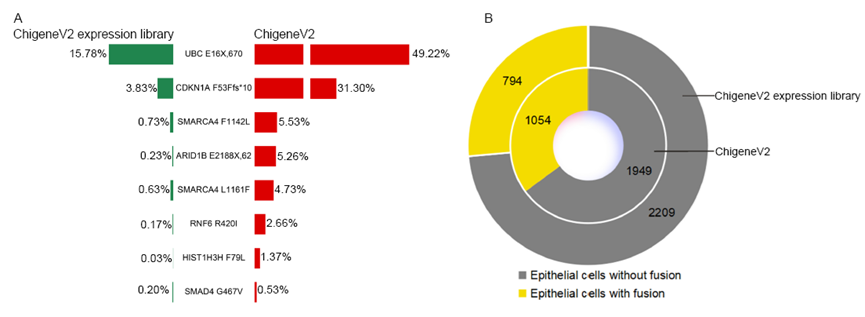

**Figure S6: Comparison of mutation and fusion detection between the ChigeneV2 expression library and ChigeneV2 platform. (A)** Comparison of Mutations Detected by ChigeneV2 Expression Library and ChigeneV2 Platform. **(B)** Distribution of Epithelial Cells with and without Fusion.
