## Supplementary methods for "Identification of Multi-landscape and Cell Interactions in the Tumor Microenvironment through High-Coverage Single-Cell Sequencing"

**Experiment Procedure for Chigene V1**

**Date: February 20, 2022**

**(Syringe Pump Version)**

### Part I: Single-Cell Capture and cDNA Synthesis

### I. Reagents Required

| **Name** | **Storage Conditions** | **Name** | **Storage Conditions** |
| --- | --- | --- | --- |
| Chigene V1 Chip | 4℃ | 5×RT Buffer | -20℃ |
| Chigene V1 Beads | 4℃ | Maxima H RTase | -20℃ |
| NLC BUFFER | 4℃ | dNTP mix（10mM） | -20℃ |
| 20×SSC | 4℃ | 1M DTT | -20℃ |
| RT enhancer 1 | 4℃ | Wash Solution | -20℃ |
| RT enhancer 2 | 4℃ | RNase inhibitor | -20℃ |
| Wash Solution A | RT | CDS-TS（30uM） | -20℃ |
| Wash Solution B | RT | EXO I Buffer | -20℃ |
| PBS | RT | EXO I | -20℃ |
| NF Water | RT | 2×KAPA mix | -20℃ |
| LowTE | RT | CDS-Primer（10uM） | -20℃ |
| Tween-20 | RT | Chip Wash Buffer | -20℃ |
| Qubit Reagent | 4℃ | DNA Purification Magnetic Beads | 4℃ |

### II. Instruments and Consumables

- Nexcelon Cell Countera
- Syringe Pump
- Microscope
- Microplate Shaker
- Hybridization Oven
- Rotating Mixer, Mini Centrifuge
- Cold Centrifuge
- PCR Machine, Qubit, Qsep 400
- Cell Counting Slide, 1.5mL Centrifuge Tubes, 2mL Centrifuge Tubes, 0.5mL Centrifuge Tubes, PCR Tubes

**III. Preparation of Reagents Before Experiment**

- **PBSI:** Prepare PBSI according to the ratio in the table below. Store at 4℃ and keep on ice when in use.

| PBSI | PBS | 1mL |
| --- | --- | --- |
|  | RNase inhibitor | 2.5uL |
|  | Total | 1002.5uL |

- **Rinse Solution:** Prepare 600uL of rinse solution for each sample, prepare immediately before use, and keep on ice.

| Rinse Solution | 20×SSC | 180uL |
| --- | --- | --- |
|  | PBSI | 402uL |
|  | 1M DTT | 6uL |
|  | 5% TWEEN-20 | 12uL |
|  | Total | 600uL |

- **Chigene V1 Lysis Solution:** Add 25uL of 1M DTT to 1mL of NLC BUFFER, mix well, and keep on ice after preparation.
- **Chip Wash Buffer:** Thaw and mix the chip wash buffer, keep ready for use.

### IV. Pre-treatment of the Chip

- Attach a 50mL syringe tightly to the syringe pump, set the syringe range to 50mL; set the injection mode to draw first then infuse; infusion volume 0.4mL, flow rate 20mL/min; draw volume 0.22mL, flow rate 10mL/min; pause 10sec.
- Expel the liquid inside the chip, draw approximately 100uL of chip wash buffer into the chip according to the above range, let it stand at room temperature for 10 minutes.
- Inject air to expel the liquid.
- Set the infusion volume to 0.42mL, flow rate 20mL/min; draw volume 0.28mL, flow rate 10mL/min, draw approximately 200uL of pre-cooled PBSI into the chip.
- Observe under a microscope for any impurities; if impurities are present, continue to inject pre-cooled PBSI once to flush until the chip is clean.

**Attention:** When injecting the chip wash buffer, do it slowly to avoid generating too many bubbles.

### V. Cell Counting and Viability Calculation

- Mix 20uL of cell suspension with 20uL of AOPI dye.
- Vortex at least 10 times to mix well.
- Remove the protective film from the top and bottom of the counting slide, take 20uL of the mixed sample, and load it into the slide.
- Select the AOPI-LiveDead mode in the software system for counting.
- Obtain the cell count and viability.

### VI. Cell Suspension Preparation

- Load 150,000 cells per chip, calculate the required cell volume X based on the cell concentration, if the volume is too small, dilute the cells, prepare 500uL of cell suspension, example as follows:：

| **Volume of Cell Suspension** | **Cells (150,000 cells)** | **Wash Solution** | **PBSI** |
| --- | --- | --- | --- |
| 500uL | X uL | 50 uL | 450-X uL |

**Attention:** After loading the cells onto the chip, there will be about 400uL of remaining cells, which can be used for other cell experiments.

### VII. Bead Suspension Preparation

- Resuspend V1 beads, take 12.5uL of beads, supplement with 72.5uL of PBSI, the bead concentration is approximately 2.4e5/ml.

### VIII. Loading Cells

- Set the injection mode to draw first then infuse; infusion volume 0.76mL, flow rate 20mL/min; draw volume 0.58mL, flow rate 10mL/min; pause 10sec.
- Draw approximately 500uL of the prepared cell suspension according to the above range, inject it into the chip, approximately 1~10 seconds.
- Observe the number of single-cell holes under the microscope.

**Key Steps:**

- Divide the chip into 3 parts, take 4 squares for cell counting in each part, and calculate the average.
- Set the infusion volume to 0.42mL, flow rate 20mL/min; draw volume 0.28mL, flow rate 10mL/min; pause 10sec, draw approximately 200uL of PBSI into the chip to flush out the cells outside the holes.
- Immediately proceed to load the beads.

### IX. Loading Beads

- Set the injection mode to draw first then infuse; infusion volume 0.34mL, flow rate 20mL/min; draw volume 0.18mL, flow rate 10mL/min; pause 10sec.
- Draw approximately 85uL of the gently resuspended beads according to the above range, and inject it into the chip, room temperature micro-shaking for 5~10 minutes.
- Observe under the microscope, try to have beads in every hole, the requirement is that the bead coverage rate is above 80%.
- Set the infusion volume to 0.42mL, flow rate 20mL/min; draw volume 0.28mL, flow rate 10mL/min, draw approximately 200uL of PBSI into the chip, flush out the beads outside the holes, this step can be repeated 3-5 times.
- Inject air into the chip to expel all the liquid in the chip. Proceed immediately with the cell lysis process. If it cannot be done immediately, place it briefly on ice.

### X. Cell Lysis and Bead Recovery

- Set the injection mode to draw first then infuse; infusion volume 0.4mL, flow rate 0.3mL/min; draw volume 0.22mL, flow rate 10mL/min; pause 10sec.
- Draw approximately 100uL of V1 lysis solution according to the above range, inject it into the chip, approximately 2 minutes, incubate at room temperature for 2 minutes.
- Set the infusion volume to 0.42mL, flow rate 20mL/min; draw volume 0.28mL, flow rate 10mL/min.
- Draw approximately 200uL of V1 lysis solution and inject it into the chip to flush out the liquid and recover it into a 1.5mL centrifuge tube.
- Set the injection mode to continuous; infusion volume 2mL, flow rate 85mL/min; draw volume 2mL, flow rate 85mL/min, repeatedly aspirate and recover.
- Observe the residual beads in the chip under the microscope, the recovery rate of the beads should be at least 75%.

### XI. Bead Cleaning

- Centrifuge the recovered beads using a benchtop centrifuge for 30 seconds, discard the supernatant.
- Add 150uL of pre-chilled wash solution, gently vortex to resuspend, centrifuge for 30 seconds, and discard the supernatant.
- Repeat step 2 three times, ensuring to completely discard the supernatant after the final wash.

### XII. Reverse Transcription

**Reverse Transcription System**

| **Component** | **Volume (uL)** |
| --- | --- |
| 5×RT Buffer | 10 |
| NF Water | 11.5 |
| dNTP mix（10mM） | 2.5 |
| MAXIAM H RTase | 2.5 |
| Rnase inhibitor | 0.5 |
| RT enhancer1 | 10 |
| CDS-TS（30uM） | 3 |
| RT enhancer2 | 10 |
| Total | 50 |

- Prepare the reverse transcription system as outlined above.
- Add 50uL of the reverse transcription system, gently vortex to mix (set vortex to the lowest speed), and resuspend the beads.
- Place on a rotating mixer, incubate at room temperature for 30 minutes. Then place the system in a hybridization oven, rotate at maximum speed, at 42°C for 60 minutes.
- After reverse transcription, centrifuge briefly for 30 seconds, discard the supernatant.
- Add 50uL of wash solution A, vortex to mix, centrifuge briefly for 30 seconds, and discard the supernatant.
- Add 150uL of wash solution B, vortex to mix, centrifuge briefly for 30 seconds, and discard the supernatant.
- Repeat step 6 once.
- After this step, the process can be paused, resuspend in wash solution B, and store at 4°C for later use.

### XIII. Exonuclease Digestion

**Digestion System**

| **Component** | **Volume (uL)** |
| --- | --- |
| EXO I Buffer | 5 |
| EXO I | 4 |
| NF Water | 41 |
| Total | 50 |

- Prepare the 50uL digestion system as outlined above.
- Centrifuge the reverse transcription product briefly, discard the supernatant, and centrifuge again to completely remove the supernatant.
- Add 50uL of the digestion system, vortex to mix, resuspend the beads.
- Place in a hybridization oven, set to maximum speed, at 37°C for 60 minutes (vortex every 10-15 minutes during this period), do not let this step proceed overnight.
- After digestion, centrifuge briefly for 30 seconds, discard the supernatant.
- Add 50uL of wash solution A, vortex to mix, centrifuge briefly for 30 seconds, and discard the supernatant.
- Add 150uL of wash solution B, vortex to mix, centrifuge briefly for 30 seconds, and discard the supernatant.
- Repeat step 7 once.
- After this step, the process can be paused, resuspend in wash solution B, and store at 4°C for later use.

### XIV. Pre-amplification

**Amplification System**

| **Component** | **Volume (uL)** |
| --- | --- |
| 2xKAPA Mix | 100 |
| CDS-Primer（10uM） | 32 |
| W3-2F（10um） | 32 |
| NF Water | 36 |
| Total | 200 |

- Centrifuge the digestion product to completely remove the supernatant, add 200uL of the amplification system, pipette to mix, and aliquot into 8 tubes for amplification (25uL per tube), with the following amplification program:

| 95℃ | 3min |  |
| --- | --- | --- |
| 95℃ | 20s | 2cycle |
| 65℃ | 20s |  |
| 72℃ | 3min |  |
| 95℃ | 20s | 13cycle |
| 65℃ | 20s |  |
| 72℃ | 30s |  |
| 72℃ | 5min |  |

**Key Steps:** For the first 2 cycles, after each denaturation step, remove the PCR tube, quickly vortex, then quickly place it back, and close the PCR machine lid.

- After the reaction, take the supernatant to measure with Qubit.
- Combine the 8 tubes of the system, add 0.6x Yeasen magnetic beads (120uL), incubate at room temperature for 5 minutes, then place on a magnetic stand until the liquid is clear, discard the supernatant, wash twice with 200uL of 80% ethanol, and then air dry at room temperature.
- Add 30uL of low-TE to resuspend the magnetic beads, incubate at room temperature for 5 minutes, then place on a magnetic stand, collect the supernatant.
- Measure with Qubit, and perform capillary electrophoresis to quality check the fragment size.
